# PTBP1/HNRNP I controls intestinal epithelial cell regeneration by maintaining stem cell survival and stemness

**DOI:** 10.1101/2021.11.03.466345

**Authors:** Wesley S. Tung, Ullas Valiya Chembazhi, Jing Yang, Ka Lam Nguyen, Aryan Lalwani, Sushant Bangru, Danielle Yee, Kristy Chin, Auinash Kalsotra, Wenyan Mei

## Abstract

Properly controlled intestinal epithelial cell regeneration is not only vital for protection against insults from environmental hazards but also crucial for preventing intestinal cancer. Intestinal stem cells located in the crypt region provide the driving force for epithelial regeneration, and thus their survival and death must be precisely regulated. We show here that polypyrimidine tract binding protein 1 (PTBP1, also called heterogeneous nuclear ribonucleoprotein I, or HNRNP I), an RNA-binding protein that post-transcriptionally regulates gene expression, is critical for intestinal stem cell survival and stemness. Mechanistically, we show that PTBP1 inhibits the expression of PHLDA3, an AKT repressor, and thereby maintains AKT activity in the intestinal stem cell compartment to promote stem cell survival and proliferation. Furthermore, we show that PTBP1 inhibits the expression of PTBP2, a paralog of PTBP1 that is known to induce neuron differentiation, through repressing inclusion of alternative exon 10 to *Ptbp2* transcript. Loss of PTBP1 results in a significant upregulation of PTBP2, which is accompanied by splicing changes in genes that are important for neuronal cell development. This finding suggests that PTBP1 prevents aberrant differentiation of intestinal stem cells into neuronal cells through inhibiting PTBP2. Our results thus reveal a novel mechanism whereby PTBP1 maintains intestinal stem cell survival and stemness through the control of gene function post-transcriptionally.

## Introduction

Intestinal epithelium lines the gastrointestinal tract and plays a critical role in digestion, absorption, and protection against insults from luminal contents. Intestinal epithelium undergoes rapid turnover driven by intestinal stem cells (ISCs) located in the crypt of Lieberkühn. ISCs continuously divide to generate new stem cells and proliferating transit-amplifying cells. Most proliferating transit-amplifying cells migrate up along the crypt-villus axis, during which they differentiate into specialized intestinal epithelial cell (IEC) lineages that execute specific physiological functions. Once the mature IECs reach the tips of villi, they undergo apoptosis and are replaced by new epithelial cells generated by ISCs. A small number of transit-amplifying cells migrate down to the crypt base and differentiate into Paneth cells which play a critical role in supporting ISCs and contributing to epithelial regeneration in response to inflammation [1–4].This self-renewal process takes approximately 3 to 5 days and is critical for maintaining intestinal epithelial integrity physiologically as well as following tissue damage [5, 6]. Based on the localization and functional properties, two major types of ISC populations have been identified. The first population is the active cycling crypt base columnar (CBC) stem cells, which are required for self-renewal of IECs under homeostatic conditions [6, 7]. The second population is the “reserve stem cells” (RSCs), which are activated in response to epithelial damage to replenish the pool of CBC stem cells and repopulate IECs [6, 7]. Given the critical role of ISCs in intestinal homeostasis, their survival and regeneration must be tightly regulated. Excessive ISC activity is a driving force of gastrointestinal cancer.

The phosphoinositide-3-kinase–Akt (PI3K-AKT) pathway is reported to be essential for maintaining the survival and regenerative capacity of ISCs under homeostatic conditions and post-injury [8–11]. AKT is a key effector of the PI3K-AKT pathway. In response to stimulation by a variety growth factors, AKT is recruited to the cell membrane via interactions between its pleckstrin homology (PH) domain and the membrane lipid phosphatidylinositol-3,4,5-trisphosphate (PIP_3_) [12]. The relocation of AKT to the cell membrane allows AKT to be phosphorylated on Thr308 and Ser473 and become activated to induce downstream signaling cascades that inhibit cell apoptosis and promote cell proliferation [12]. While AKT is critical for ISC-mediated epithelial cell regeneration, excessive AKT activity induces intestinal tumorigenesis as indicated by frequent activating mutations in the PI3K-AKT pathways in human colorectal carcinomas [13]. These observations highlight the importance of well-balanced PI3K-AKT signaling activity in intestinal homeostasis.

The activity of PI3K-AKT signaling is reported to be antagonized by tumor suppressor gene *p53* in several tissues and cell lines [14–19]. One of the P53-mediated AKT inhibitory mechanisms is through the transcriptional activation of AKT repressors. PH-like domain family A, member 3 (PHLDA3) is one such repressor induced by P53, and it possesses a PH domain that allows it to act as a dominant-negative form of AKT [20, 21]. In cultured cells, PHLDA3 prevents AKT activation via interfering with AKT binding to PIP_3,_ resulting in apoptosis [20, 21]. While it remains unclear whether PHLDA3 functions *in vivo* to regulate the AKT activity in the ISC compartment, frequent mutations and copy number variation of *Phlda3* gene were found in colon carcinoma (https://cancer.sanger.ac.uk/cosmic), which highlights a potential role of PHLDA3 in regulating intestinal homeostasis.

Polypyrimidine tract binding protein 1 (PTBP1, also called HNRNP I) is an RNA-binding protein that serves critical roles in post-transcriptional gene regulation, particularly as a repressor of exon inclusion [22–25]. Increasing evidence indicates that PTBP1-mediated post transcriptional regulation is essential for a number of biological processes, ranging from embryonic development, neuronal cell differentiation, immune development, intestinal homeostasis, to spermatogenesis [26–34]. Yet, how PTBP1 executes its physiological functions in a tissue-specific manner remains poorly understood. We previously reported that deletion of *Ptbp1* gene in mouse neonatal IECs disrupts neonatal immune adaptation and causes early onset of colitis and colorectal cancer [34]. Here, we report that in adulthood, PTBP1 controls the survival and death of ISCs and the transit-amplifying cells, and is indispensable for IEC regeneration. This is at least in part through balancing the activities of AKT and P53. Mechanistically, this occurs due to PTBP1-mediated repression of PHLDA3 expression in the ISC compartment, which provides a permissible environment for AKT activation. We further show that PTBP1 inhibits its paralog PTBP2 in crypt cells and controls splicing programs important for neuron differentiation, which in turn maintains the undifferentiated states of ISCs and the transit-amplifying cells. Our results thus reveal a novel mechanism whereby PTBP1 controls ISC survival and stemness through regulating gene function post transcriptionally.

## Results

### IEC-specific *Ptbp1* deletion in adulthood results in impaired intestinal epithelium regeneration

Previously, we investigated the function of PTBP1 in the intestinal epithelium by deleting the *Ptbp1* gene in IECs from early embryogenesis using *Villin*-Cre [34]. While this mouse model allowed us to gain valuable insights into the role of PTBP1 in neonatal IECs, it is not applicable to study the function of PTBP1 in the adult intestinal epithelium. Therefore, we generated a tamoxifen-inducible *Ptbp1* knockout mouse model *Ptbp1*^f/f^; *Vil-cre*^ER+/−^ mice (See Materials and methods). Assessment of *Ptbp1* mRNA and protein expression in knockout mice 48 hours post tamoxifen induction (PTI) revealed near-complete depletion of PTBP1 protein in the IECs (Fig. 1D to 1F).

**Figure 1.**
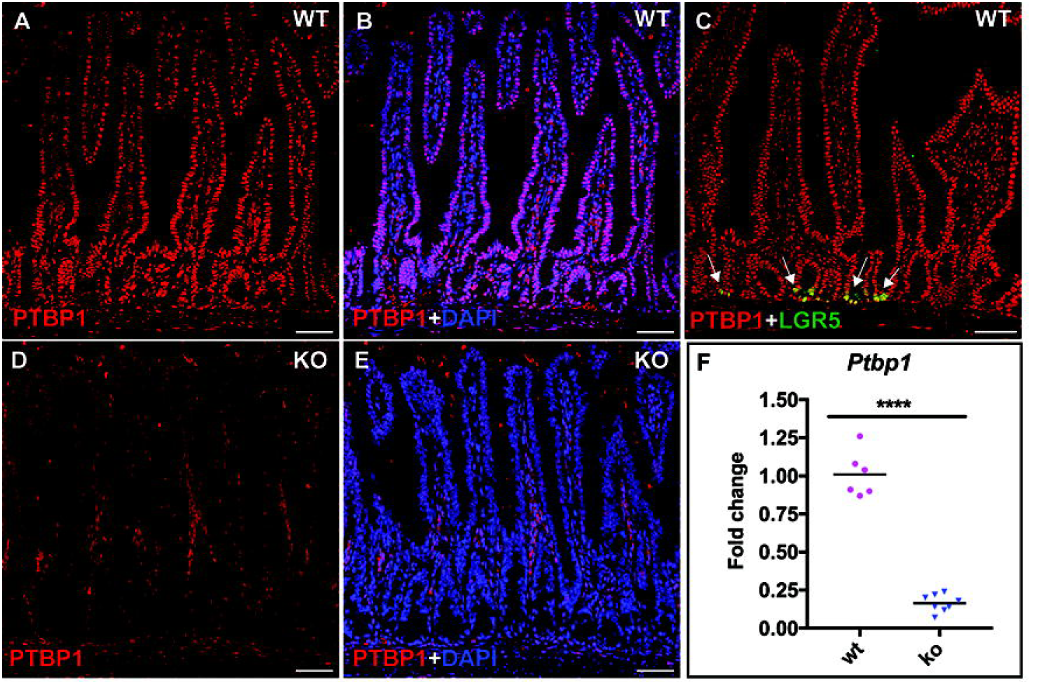
Generation of the IEC-specific *Pthpl* knockout mice. (A-E) Immunohistochemical staining using an anti-PTBPl antibody shows PTBPl protein localization in the wild-type and *Ptbpl* knockout small intestine. Nuclear accumulation of PTBPl protein was detected in all IECs including LGR5-positive CBC stem cells (arrows in C) in wild-type mice. LGR5-posititve CBC stem cells were detected by an anti-GFP antibody on intestinal tissues from *Ptbpl*^*f/f*^*;Lgr5* mice. PTBPl expression is diminished in the IECs in the knockout mice, but remains unaffected in the lamina propria cells at 48 hours PTI (D). Nuclei were counterstained with DAPI. (E) Quantitative Real-time PCR shows significant decreased level of *PtbplmRNA* in the intestinal crypt cells of the knock-out mice at 48 hours PTI. Data were normalized to *Gapdh*. Each symbol in the graphs indicates fold change of *Ptbpl* expression in individual mice. Bars show mean values. N **=** 8 knockout mice and 6 control mice. Mice of the knockout group and the wild-type group are sibling littermates. Data shown are representative of at least three independent experiments. **** p **<** 0.0001. WT, wild-type; KO, knockout. Scale bars, 50 μm.

Strikingly, all *Ptbp1* knockout mice showed significant weight loss and died within 7 days PTI (Fig. 2A and 2B). To determine the cause of death, we collected intestine tissues every 24 hours PTI and performed histological analysis. We found that while the overall organization of intestinal epithelium in the knockout mice appeared relatively normal at 24 and 48 hours PTI, a small number of apoptotic cells appeared in the crypt region starting at 24 hours PTI (data not shown), and the number of apoptotic cells dramatically increased at 48 hours PTI (Fig. 2D, Table 1). By 72 hours PTI, knockout mice displayed severe crypt death, accompanied by infiltration of immune cells in the lamina propria (Fig. 2F). The regular crypt-villus architecture was completely disrupted in the knockout mice at 96 hours PTI, displaying as villous atrophy (Fig. 2H). Cell apoptosis in the crypt region of the knockout mice was confirmed by a dramatic increase in the number of cleaved CASPASE3-positive cells in the knockout mice at 48 hours PTI (compare Fig. 3B to 3A, Table 1). While crypt cell death was observed in both the small and large intestines in the knockout mice, the small intestine was more severely affected by PTBP1 deletion.

**Figure 2.**
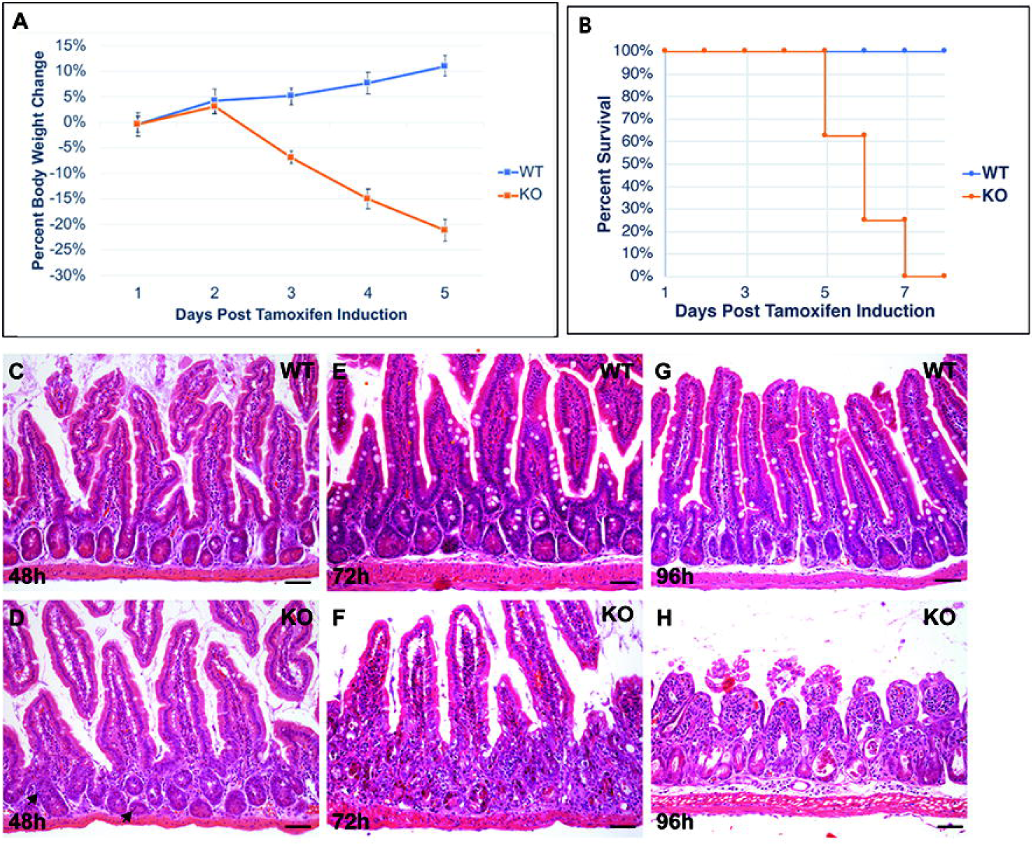
Loss of epithelial PTBPl results in failure in IEC regeneration. (A) shows weight loss of *Ptbpl* knockout mice within 5 days PTI. (B) shows death of *Ptbpl* knockout mice within 7 days PTI. Data in A and B were from N = 8 knockout mice and 9 control mice. (C-H) Hematoxylin and Eosin-stained sections show changes of the epithelial structure at 48 hours (C and 0), 72 hours (E and F), and 96 hours (G and H) PTI in the control mice and *Ptbpl* knockout mice. Note that the knockout mice displayed villi atrophy at 96 hours PTI. Arrows in D show apoptotic cells. WT, wild-type; KO, knockout. Scale bars, 50 μm.

**Table 1:**
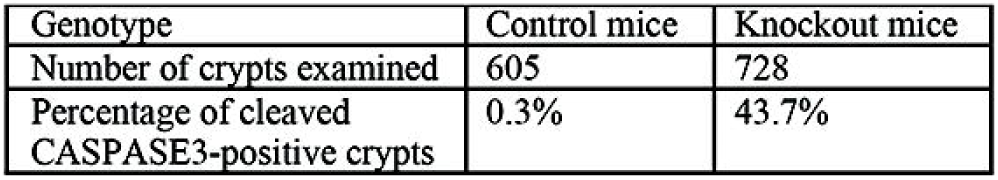
Quantification of crypt cell death

**Figure 3.**
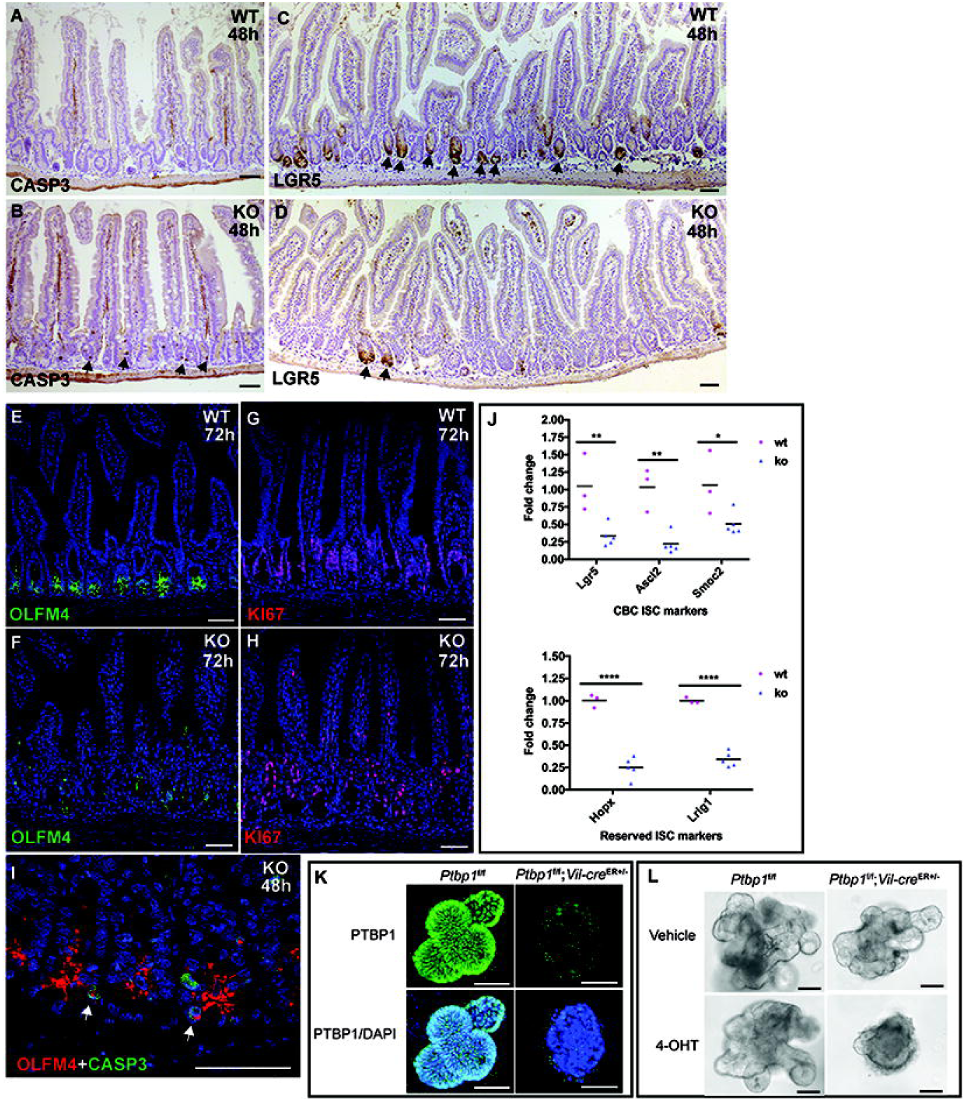
PTBPl deficiency results in loss of intestinal stem cells. (A-B) lmmunohistochemical staining with an anti-cleaved CASPASE 3 antibody shows increased number of apoptotic cells in the crypt region of knockout mice at 48 hours PTI (8, arrows). (C-D) lmmunohistochemical staining with an anti-GFP antibody shows LGR5-postitive CBC stem cells, the number of which is significantly decreased in the knockout mice at 48 hours PTI (D). Arrows point to LGRS-posititive crypt cells. (E-H) lmmunohistochemical staining shows knockout mice have a significant decrease in the number ofOLFM4-posititve cells (F) and Kl67-postitive cells at 72 hours PTI (H). (I) Double immunohistochemical staining with anti-OLFM4 and anti-cleaved CASPASE 3 antibodies shows death ofOLFM4-posititve stem cells (arrows in I). (J) Real-time PCR shows decreased expression of CBC stem cell markers and RSC markers. Data were normalized to *Gapdh*. Each symbol in the graphs indicates fold change of gene expression in individual mice. Bars show mean values. N = 5 knockout mice and 3 control mice. Mice of the knockout group and the wild-type group are sibling littermates. All data shown are representative of at least three independent experiments.* p < 0.05; ** p < 0.01; **** p < 0.000 I. (K) Whole mount immunostaining with an anti-PTBPl antibody shows efficient depletion of PTBPlin 4-OHT-treated organoids derived from *Ptbpl*^f/f^;*Vil-cre*^ER+/−^. mice but not those from *Ptbpl*^f/f^ mice at day 3 of culture. Organoids were counterstained with DAPI. (L) 4-OHT-treated organoids derived from *Ptbpl*^f/f^;*Vil-cre*^ER+/−^ mice failed to bud at day 4 of culture, but not those from the control mice. Vehicle-treated organoids from *Ptbpl*^f/f^;*Vil-cre*^ER+/−^ mice and the control mice displayed nonnal budding. WT, wild-type; KO, knockout. 4-OHT, hydroxytamoxifen. Scale bars, 50 μm.

### Loss of ISCs in *Ptbp1* knockout mice

Since ISCs fuel the intestinal epithelial regeneration, we checked if villous atrophy in the knockout mice is caused by the loss of ISCs. We generated a compound *Ptbp1* knockout mouse model *Ptbp1*^f/f^;*Vil-cre*^ER+/−^;*Lgr5* in which LGR5-positive CBC stem cells are labeled with GFP (see Materials and methods). We found that the number of LGR5-positive ISCs is significantly reduced in the knockout mice at 48 hours PTI (Fig. 3D, Table 2). Consistent with this finding, we found expression of OLFM4, another CBC stem cell marker [35], is diminished in the crypt cells of the knockout mice at 72 hours PTI (Compare Fig. 3E to 3F). Furthermore, we detected cleaved CASPASE3 staining in OLFM4-positive ISCs at 48 hours PTI (Fig. 3I), suggesting that loss of ISCs is due to apoptosis of these cells in the knockout mice. To determine whether both CBC stem cells and RSCs are affected, we performed qPCR to assess the expression level of genes specific for CBC stem cells or RSCs. We found that the expression levels of *Lgr5, Ascl2*, and *Smoc2* (signature genes for CBC stem cells), and *Hopx* and *Lrig1* (RSCs markers) were significantly reduced in the knockout mice (Fig. 3J), indicating loss of both ISC populations. In support of this finding, we detected a significant reduction in the number of proliferating transit-amplifying cells at 72 hours PTI in the knockout mice (compare Fig. 3G to 3H). These results indicate that the epithelial PTBP1 plays a critical role in maintaining the survival and proliferation of ISCs.

**Table 2:** Quantification of LGRS;EGFP-positive crypts

| Genotype | Control mice | Knockout mouse 1 | Knockout mouse 2 |
| --- | --- | --- | --- |
| Number of crypts examined | 848 | 843 | 783 |
| Percentage of LGR5;EGFP-positive crypts | 38.0% | 8.9% | 9.7% |

To further determine if PTBP1 is required for maintaining the ISC regenerative capacity, we established *ex vivo* intestinal organoid cultures using crypts isolated from *Ptbp1*^f/f^; *Vil-cre*^ER+/−^ mice and their control littermates *Ptbp1*^f/f^ mice. Efficient deletion of *Ptbp1* was observed upon treating organoids with 4-hydroxytamoxifen for 24 hours (Fig. 3K). We found that 4-hydroxytamoxifen-treated *Ptbp1*^f/f^; *Vil-cre*^ER+/−^ crypts failed to bud at day 3 of culture when compared to vehicle-treated *Ptbp1*^f/f^; *Vil-cre*^ER+/−^ crypts or 4-hydroxytamoxifen-treated *Ptbp1*^f/f^ crypts (Fig. 3L). This observation indicates that PTBP1 is required for ISC-mediated epithelial regeneration.

### Transcriptome changes in PTBP1-deficient cells

To determine the molecular mechanism by which PTBP1 controls intestinal survival and regeneration, we deep-sequenced poly(A) selected RNAs prepared freshly from crypt cells of age-matched wild-type and PTBP1-deficient mice. We chose to assess the transcriptome changes in the knockout mice at 20 hours PTI (4 hours before cell death could be detected in the crypt region) so that the observed changes represent the direct consequence of PTBP1 deficiency and are likely the cause of the crypt cell death. Our results show that PTBP1 deletion predominantly affects mRNA splicing (Fig. 4A). We identified a total of 1165 altered splicing events within 834 genes (Figure 4A and 4B). Remarkably, in contrast to the large number of splicing changes, only 26 genes showed changes in their mRNA abundance at 20 hours PTI (Fig. 4A). Amongst them, 4 genes (*Trim72, Ager, H2-Bl*, and *Fmr1nb*) exhibited significant differences in both mRNA abundance and splicing. These findings provide strong evidence that the splicing defects triggered by PTBP1 deficiency are primary events that precede the onset of crypt cell death and immune cell infiltration in the PTBP1 knockout intestines.

**Figure 4.**
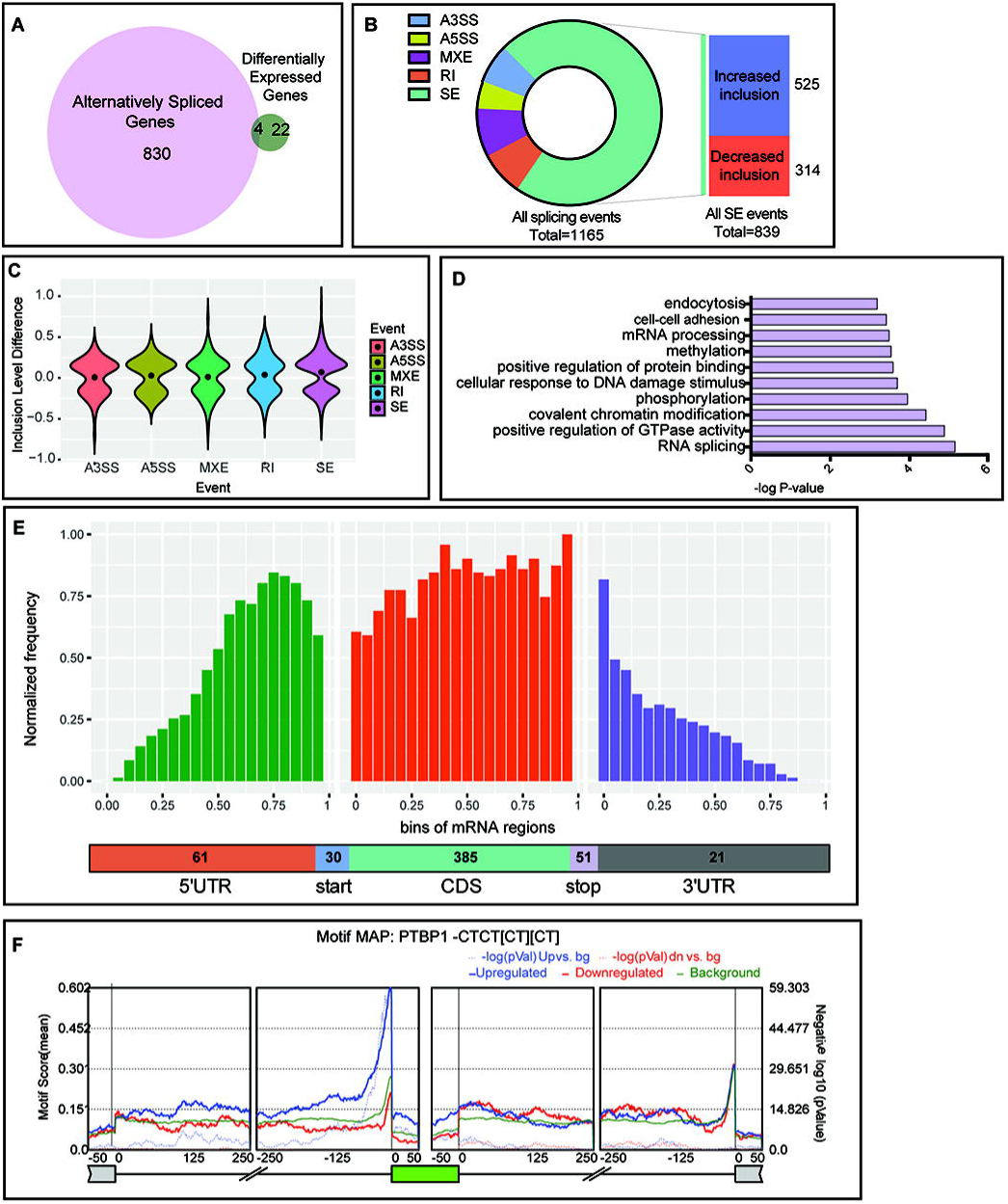
Changes of transcriptome in PTBPl-deficient crypt cells. (A) The number of genes in which differential gene expression/alternative splicing changes were detected in the knockout crypt cells. Four genes *(Trim 72, Ager, H2-Bl* and *Fmrlnb)* displayed changes in both gene expression and splicing. (B) Breakdown of 1165 alternatively spliced events into various event categories. A3SS: alternative 3’ splice site, A5SS: Alternative 5’ splice site, MXE: Mutually exclusive exon, RI: Retained intron, SE: Skipped exon. (C) Violin plots demonstrating t,.PSI(lnclusion level difference) distributions of significantly altered splicing events in the knockout crypt cells. (D) Gene ontology analysis demonstrating top 10 biological processes that are enriched among genes that are alternatively spliced. (E) Metagene analysis of alternatively spliced exons in knockout mice by their positions on an mRNA transcript. Distribution along the transcript bins is shown on top, and breakup of events into relative transcript regions is shown in the bottom. (F) shows the relative enrichment of PTBPl binding motifnear cassette exons (represented in green) that displayed significantly increase (blue curve) or decrease (red curve) in inclusion in PTBPl-deficient crypt cells. Alternatively spliced exons were identified using rMATS and motif map was constructed using RMAPs (with 50-nucleotide sliding window). The set of background cassette exons is represented in black.

The majority of splicing changes were exon skipping events (72%), but changes in alternative 5’ or 3’ splice sites, intron retention, and mutually exclusive exons were also detected (Fig. 4B). Notably, nearly two-thirds of skipped exons displayed increased inclusion in PTBP1-deficient crypt cells (Fig. 4C), which is consistent with the previous reports that PTBP1 primarily functions as a repressor of splicing [25]. Gene ontology analysis further revealed that the differentially spliced mRNAs following *Ptbp1* deletion were enriched in several functional clusters, including mRNA splicing, regulation of GTPase activity, chromatin modification, phosphorylation, and cellular response to DNA damage stimulus (Fig. 4D).

We next investigated the spatial distribution of PTBP1-regulated exons in the crypt cells along their associated transcripts. Metagene analysis revealed that approximately 70% of differentially spliced exons were located within coding sequences (CDS) and a sizable number of those (15%) encoded an alternate START or STOP codon (Fig. 4E). Furthermore, we found a strong overrepresentation of PTBP1 binding motif (CUCUCUCU) near the 3’ splice site of the upstream introns surrounding the alternative exons that were abnormally included in PTBP1-deficient crypt cells (Figure 4F). This suggests that in crypt cells, PTBP1 suppresses the inclusion of many alternate exons by directly binding to its motif in the upstream introns.

The CDS-mapped PTBP1-regulated exons were further classified based on whether they were open reading frame preserving (exon length is a multiple of 3). We found that 70% of the regulated exons preserve the open reading frame, whereas 30% of them do not (Figure 5A). In both cases, only about 3% of the exons were predicted to undergo nonsense-mediated RNA decay (Fig. 5A). This result is consistent with our transcriptome data wherein most of the mRNAs harboring PTBP1-regulated exons in crypt cells do not exhibit a significant change in their overall abundance (Fig. 4A). On the contrary, these exons are likely to alter the intrinsic structure and function of the encoded proteins. To further probe the functional properties and features of PTBP1-regulated exons, we performed exon ontology analysis, which revealed a significant enrichment for sequences encoding intrinsically unstructured regions, phosphorylation sites, post-translational modifications, cellular localization, binding, and catalytic activity (Fig. 5B and 5C). We further noted that within the cellular localization category, many PTBP1-regulated exons contained a nuclear localization, export, or a membrane targeting signal (Fig. 5D).

**Figure 5.**
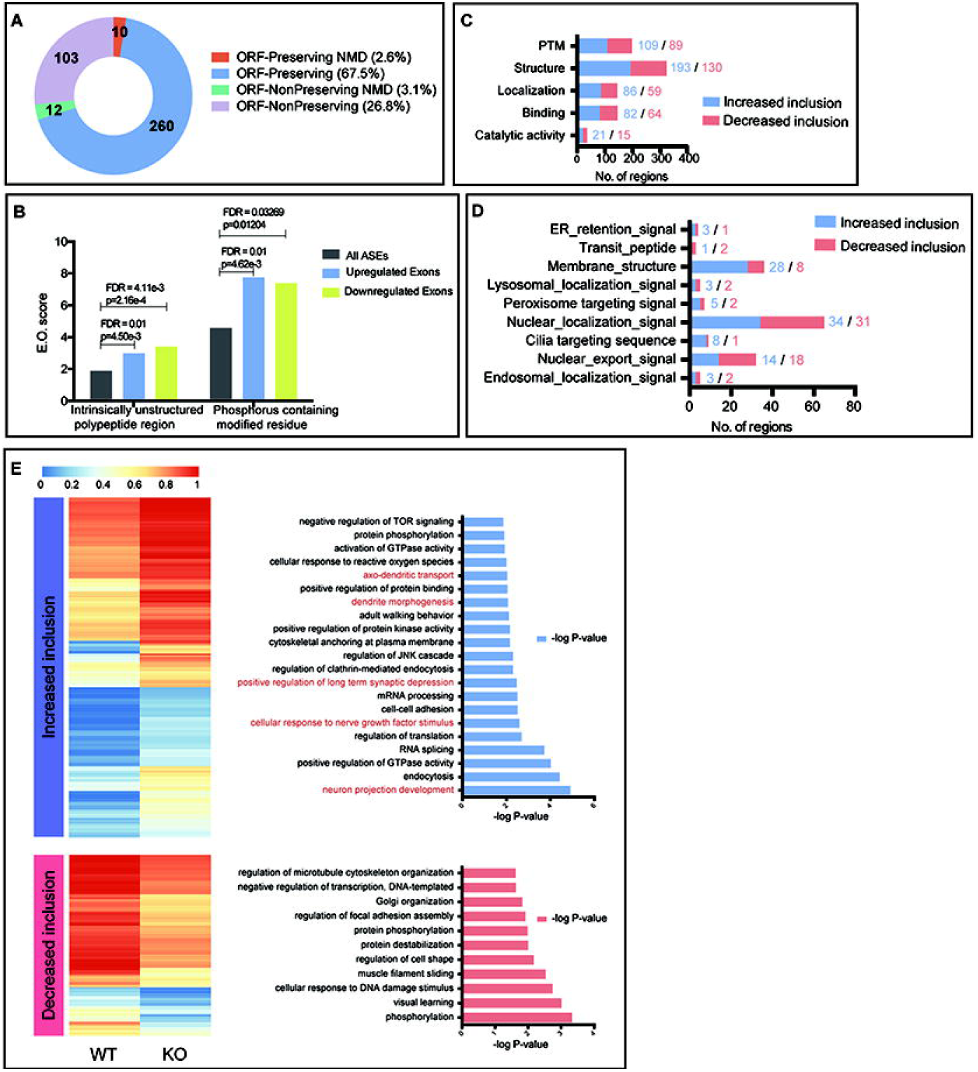
Detailed analysis of exons in the skipped exon category. (A) Effect of alternatively spliced events induced by PTBP I deficiency on the transcripts in crypt cells. Categories indicate if the exon inclusion preserved the open reading frame (ORF) and if the inclusion transcript is subjected to Non-sense Mediated Decay (NMD). (B) Relative to a set of all Alternatively Spliced Exons (ASEs), PTBPl-regulated exons (either upregulated or downregulated) are significantly enriched for regions encoding intrinsically unstructured regions and phosphorylation sites. Exon Ontology(E.O.) scores are shown on they axis. (C) Exon ontology-based distribution of skipped exons, either increase or decrease in inclusion, based on encoded protein features. PTM: post translational modification. (D) Exon ontology-based distribution of differentially spliced exons based on localization. (E) Heatmap showing average PSI values of exons that have either significantly increased (top) or decreased (bottom) inclusion in the knockout mice. Corresponding gene ontology analysis reveals biological process that are enriched among these genes (shown on the right). Neuron cell differentiation-related biological processes are highlighted in red.

### PTBP1 suppresses PTBP2 expression at the mRNA and protein levels in crypt cells

Gene ontology analysis revealed that the differentially spliced genes due to altered exon skipping in PTBP1-deficient crypt cells function in a number of biological processes, ranging from mRNA processing and translation, protein phosphorylation, cellular morphogenesis, to signal transduction (Fig. 5E). Interestingly, we found that 5 of these biological processes were related to neuronal cell differentiation, including neuron projection development, cellular responses to nerve growth factor stimulus, regulation of synaptic depression, dendrite morphogenesis, and axo-dendritic transport (Fig. 5E). This finding prompted us to check the expression pattern of *Ptbp2* (also called neuronal *Ptb*), a paralog of *Ptbp1* that is highly expressed in the neuronal cells and is required to initiate the neuron-specific splicing program [36–38]. PTBP1 is known to inhibit *Ptbp2* expression in neuronal progenitor cells and many non-neuronal cells in the mouse brain by repressing the inclusion of the alternative exon 10 in the *Ptbp2* transcript, which leads to nonsense-mediated RNA decay of *Ptbp2* [28, 36]. To determine if a similar repression mechanism exists in the crypt cells, we assessed exon 10 inclusion in *Ptbp2* mRNA. Indeed, we observed a significant increase in the inclusion of *Ptbp2* exon 10 in PTBP1-deficient crypt cells at 20 hours PTI (Fig. 6A to 6C). This was accompanied by a dramatic upregulation of *Ptbp2* expression at both mRNA and protein levels (Fig. 6D and 6E). These results strongly suggest that PTBP1 inhibits PTBP2 expression in the ISC compartment and serves a critical role in suppressing the neuron-specific splicing program in ISCs.

**Figure 6.**
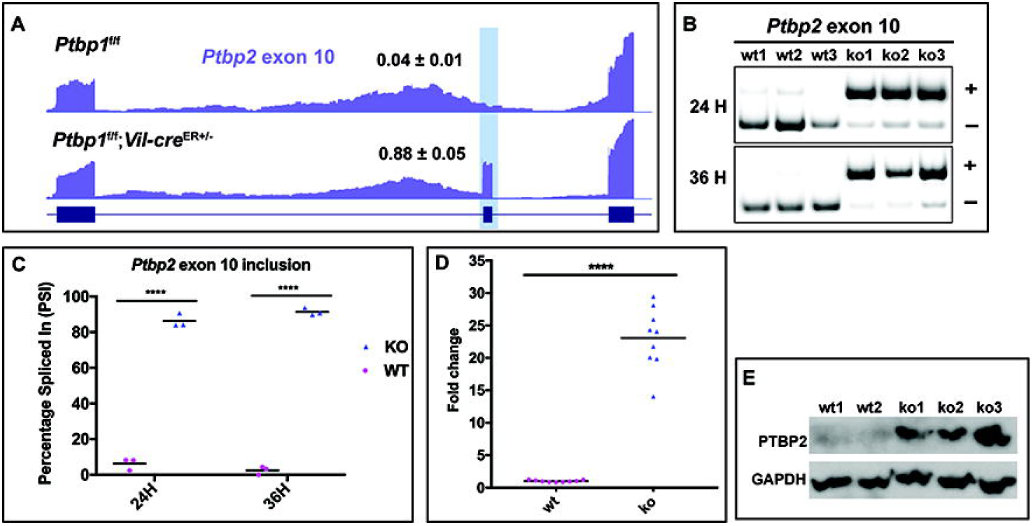
Loss of PTBPl increased *Ptbp2* expression both at the mRNA and protein levels. (A) UCSC Track image shows high inclusion of exon 10 of *Ptbp2* in the knockout crypt cells. (B-C) PCR-based splicing assay on exon 10 inclusion in *Ptbp2* mRNAat 24 and 36 hours PTI. Gel images show a significant increase in exon 10 inclusion (B). Exon inclusion and exclusion bands are denoted by(+) and(-) respectively. Quantification was done using Image Lab 5.2.1 software (Biorad). PSI values were determined using lmageLab software (BioRad) as the exon inclusion band intensity/(the exon inclusion band intensity+ the exon exclusion band intensity) x 100. Bars show mean values (C). N = 3 WT and 3 KO mice at each time point. WT and KO mice used for analysis are littermates. (D) Real-time PCR shows a significant upregulation of *Ptbp2* mRNA at 20 hours PTI. Data were normalized to *Gapdh*. Each symbol in the graphs indicates fold change of *Ptbp2* expression in individual mice. Bars show mean values.**** p < 0.0001. In both the wild-type and knockout groups, N = 9 mice. (E) Western blot shows PTBP2 protein expression is increased in the knockout crypt cells at 24 hours PTI. GAPDH is served as the loading control. WT, wild-type; KO, knockout. All data shown are representative ofat least three independent experiments.

### PTBP1 downregulates PHLDA3 and maintains AKT activity in the ISC compartment

Given that loss of PTBP1 causes crypt cell apoptosis, we analyzed genes associated with cell apoptosis among those that are alternatively spliced or differentially expressed upon loss of PTBP1. This led to the identification of *Phlda3*, a gene that encodes a negative regulator of AKT signaling activity. *Phlda3* showed the highest fold change among the upregulated genes in PTBP1-deficient crypt cells (Fig. 7A). We performed real-time PCR to assess the expression of *Phlda3* in PTBP1-deficient crypt cells at different time points after tamoxifen induction. An increase in *Phlda3* expression was first detected at 20 hours PTI in PTBP1-deficient crypt cells and became more prominent at 36 and 48 hours PTI (Fig. 7B). Immunostaining revealed that PHLDA3 protein was accumulated in the cells at the crypt bottom where ISCs and Paneth cells are located (Fig. 7C). In the PTBP1 knockout mice, PHLDA3 protein expression is significantly increased in the crypt bottom region (Fig. 7D). To determine which cell lineages express PHLDA3 protein, we performed double immunofluorescence staining with anti-PHLDA3 and anti-GFP antibodies on intestines of *Ptbp1*^f/f^; *Lgr5* mice in which LGR5-expressing CBC stem cells are labeled with GFP. Interestingly, we found PHLDA3 protein is localized in cells adjacent to LGR5-positive CBC stem cells (Fig. 7E). Furthermore, these PHLDA3-expressing cells contain lysozyme, indicating that they are Paneth cells (Fig. 7F).

**Figure 7.**
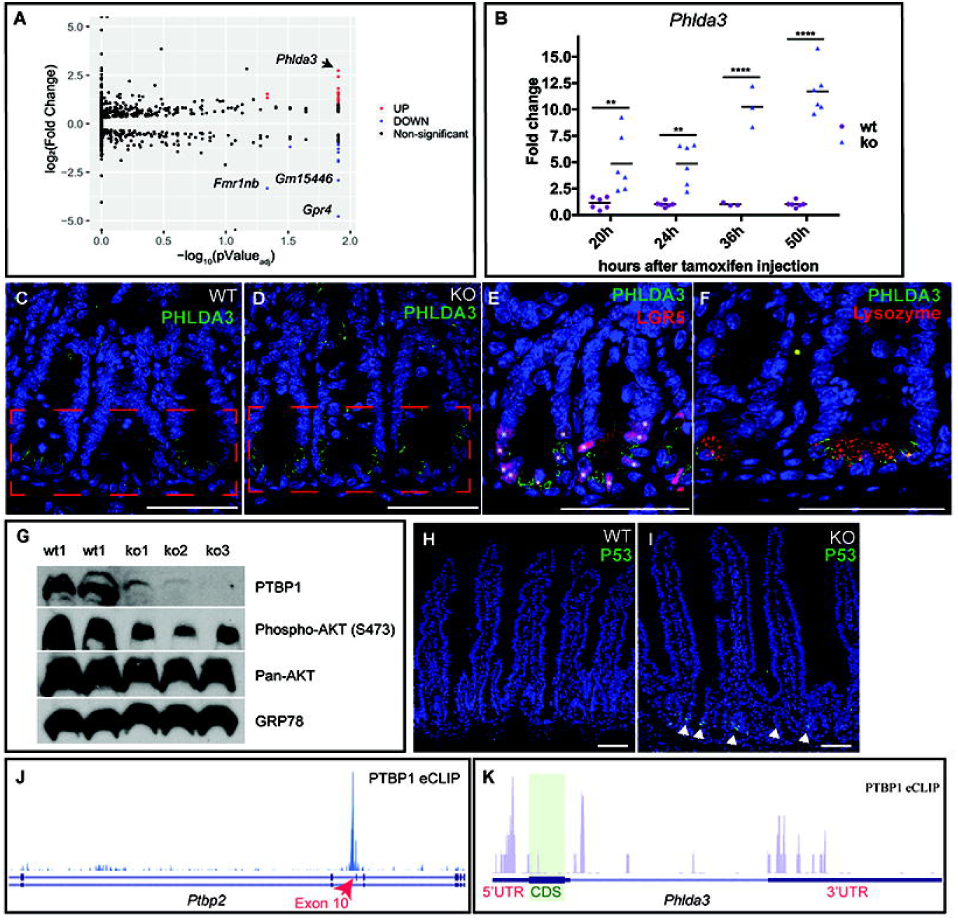
Loss of PTBPl upregulates PHLDA3 and impairs AKT activation in crypt cells. (A) Volcano plot demonstrates mRNA abundance changes in the knockout crypt cells. Red dots represent significantly upregulated genes (log2FC> 1, FDR<0.05), blue dots represent significantly downregulated genes (log2FC<-l, FDR<0.05) and the black dots represent genes with no significant changes in expression between the knockout and control crypt cells. *Phlda3* showed the highest fold change among the upregulated genes (arrow). (B) Real-time PCR shows a significant upregulation of *Phlda3* mRNA level at 20, 24, 36, and 50 hours PTI. Data were normalized to *Gapdh*. Each symbol in the graphs indicates fold change of *Phlda3* expression in individual mice. Bars show mean values. At each time point, sibling littermates were used for analysis and in both the wild-type and knockout groups, N = 6 knockout mice and 6 control mice for 20, 24, and 50 hours PTI. N = 3 knockout mice and 3 control mice for 36 hours PTI. “““ p < 0.01; **** p < 0.0001. Data shown are representative of at least three independent experiments. (C-D) lmmunohistochemical staining using an anti-PHLDA3 antibody shows PHLDA3 protein expression is increased in the bottom crypt cells in the knockout mice (D). (E) Double Immunofluorescence staining using anti-PHLDA3 and anti-GFP antibodies shows PHLDA3 protein is localized in cells adjacent to LGR5-positive CBC stem cells(*) in wild-type mice. (F) Double lmmunofluorescence staining using anti-PHLDA3 and anti-lysozyme antibodies shows PHLDA3 protein is localized in Paneth cells in wild-type mice. (G) Western blot shows decreased AKT phosphorylation on ser473 in knockout crypt cells at 24 hours PTI. 5 mice used in the assay are littennates. PTBPl was efficient depleted in all 3 knockout mice. (H-1) Immunohistochemical staining with an anti-P53 antibody shows an upregulation of P53 in the knockout crypt cells at 48 hours PTI as indicated by nuclear localization of P53 (arrows in I). (J-K) UCSC track visualization of PTBPl eCLIP data from HepG2 cells demonstrating PTBPl binding on *Ptbp2* transcript **(J)** and *Phlda3* transcript (K). eCLIP was obtained from ENCODE database. The alternative exon 10 of *Ptbp2* is indicated by red arrow in J. CDS, coding sequences. WT, wild-type; KO, knockout. Scale bars, 50 μm.

Since PHLDA3 can interfere with AKT binding to PIP_3_ and thereby prevent AKT activation [20], we thus assessed AKT phosphorylation in PTBP1-deficient crypt cells. Indeed, phosphorylation of AKT on Ser473 was significantly decreased in PTBP1-deficient crypt cells at 24 hours PTI (Fig. 7G). This was accompanied by an increase in P53 activity at 48 hours PTI, judged by nuclear localization of P53 in the crypt cells (compare Fig. 7H to 7I). Intriguingly, although the mRNA level of *Phlda3* was upregulated significantly in PTBP1-deficient crypt cells at 24 hours PTI, we did not detect any upregulation of P53 at this time point (supplemental Fig.1). This result suggests that upregulation of *Phlda3* mRNA level in PTBP1-deficient crypt cells at 24 hours PTI is induced by a mechanism independent of P53.

Finally, to investigate direct regulation of target genes including *Phlda3* by PTBP1, we analyzed the publicly available HepG2 PTBP1 eCLIP data from ENCODE database (PMID: 22955616; PMCID: PMC3439153). We detected a prominent PTBP1 eCLIP peak near the 3’ splice site in the upstream intron of *Ptbp2* exon 10 (Fig. 7J), which is known to suppress the inclusion of exon 10. While no PTBP1 binding was detected in the coding region of *Phlda3*, eCLIP tags were mapped to both 5’ and 3’ untranslated regions of the *Phlda3* transcript (Fig. 7K).

Collectively, our results reveal a novel mechanism through which PTBP1 controls ISC function post transcriptionally (Fig. 8). We demonstrate that PTBP1 downregulates PHLDA3 expression in the ISC compartment to permit AKT activation, which is essential for the maintenance of ISC survival and regeneration. We further show that PTBP1 represses PTBP2 expression in the crypt cells through the splicing control to prevent their aberrant differentiation into neurons.

**Figure 8.**
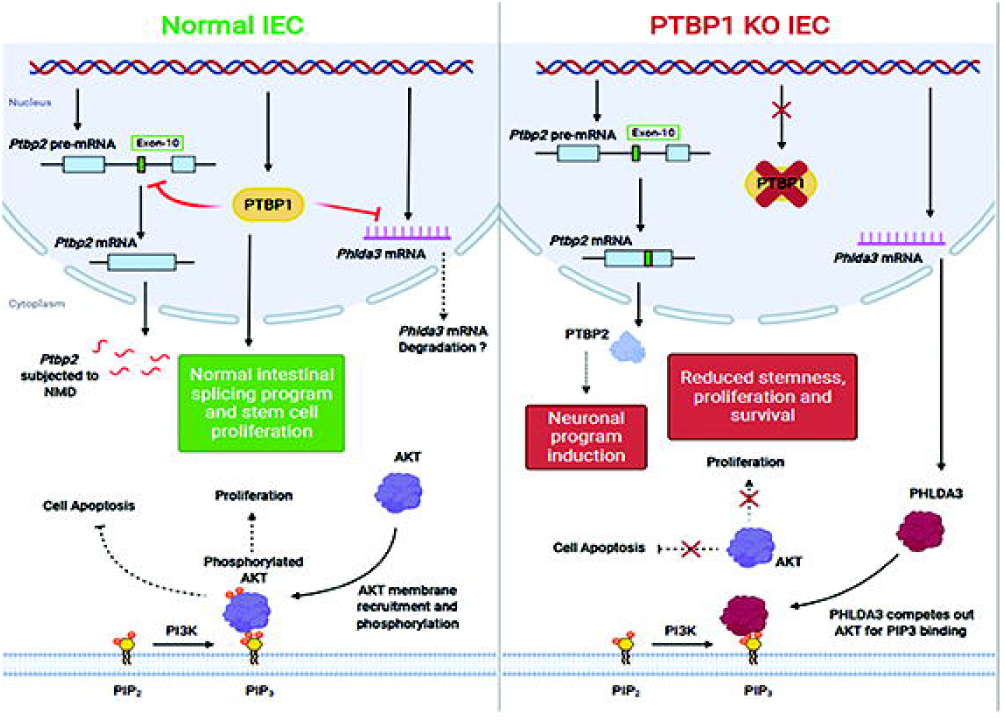
A proposed model for PTBPl function in regulating ISCs. PTBPl maintains ISC survival, proliferation, and sternness through post-transcriptionally regulating *Phlda3* and *Ptbp2* expression. PTBPl inhibits *Phlda3*, presumably through destabilizing its mRNA, which pennitsAKT phosphorylation and subsequent activationtopromotelSCsurvivalandprolifcration. PTBPl represses inclusion of *Ptbp2* exon 10, leading to nonsense-mediated mRNA decay (NMD)of *Ptbp2*. which in tum prevents induction of neuronal differentiation program to maintain ISC sternness.

## Discussion

The intestinal epithelial regeneration process is not only vital for digestion and absorption but also for protection against insults from environmental hazards. ISCs in the crypt region are the driving force of IEC renewal under both physiological conditions and upon injury. Yet, excessive ISC activity can trigger tumorigenesis. As such, the survival and regeneration of ISCs must be tightly controlled. AKT is crucial for ISC survival and proliferation [8–11], but the mechanisms that balance AKT activity in the intestine remain unclear. We show here that PTBP1, an RNA-binding protein that controls gene function post transcriptionally, is critical for regulating AKT. We demonstrate that PTBP1 maintains AKT activity through repressing the expression of PHLDA3, an AKT repressor that promotes cell death [20, 21, 39]. Loss of PTBP1 in crypt cells impairs AKT phosphorylation and results in death of ISCs and the failure of epithelial regeneration. Interestingly, we show that PHLDA3 is expressed in Paneth cells which are known to nurture and protect ISCs [1–4]. This raises a possibility that that PTBP1maintains ISC survival through the control of the niche provided by Paneth cells.

Mechanistically, we show that PTBP1may inhibit *Phlda3* expression in the ISC compartment through a direct regulation of its mRNA stability. Although PHLDA3 was previously identified as one of the P53-induced AKT repressors [20, 21], we found that the *Phlda3* mRNA level was upregulated in PTBP1-deficient crypt cells prior to detectable P53 activation, indicating that PTBP1controls *Phlda3* expression in the ISC compartment at least in part through a mechanism independent of P53. In support of this view, we identified eCLIP tags that were mapped to both 5’ and 3’ untranslated regions of the *Phlda3* transcript in the publicly available HepG2 PTBP1 eCLIP data, which suggests a direct binding of PTBP1 to the *Phlda3* transcript. Notably, PTBP1 can control the functions of its target genes either through regulating their alternative splicing or controlling their mRNA stability [22–25]; PTBP1 destabilizes mRNAs of a number of genes through direct binding to their 3’ untranslated regions [40–43]. We found *Phlda3* was differentially expressed but not alternatively spliced in PTBP1-deficient crypt cells, raising the possibility that PTBP1 destabilizes *Phlda3* mRNA in the ISC compartment through binding to its untranslated region.

We further show that PTBP1 inhibits *Ptbp2* expression in the crypt cells through repressing the inclusion of its alternative exon 10. Loss of PTBP1 results in a significant upregulation of *Ptbp2* expression at both the mRNA and protein levels. Similar mechanisms were reported in the mouse brain, where PTBP1 inhibits *Ptbp2* expression in neuronal progenitor cells and many non-neuronal cells to suppresses the splicing program that drives neuron cell differentiation [36, 44]. Consistently, loss of PTBP1 in the crypt cells induced splicing alterations in genes required for neuronal cell differentiation. This finding indicates that PTBP1 prevents aberrant differentiation of ISCs and transit-amplifying cells into neuronal cells, which is presumably critical for maintaining these cells multipotent and viable (Fig. 8). Interestingly, PTBP1 is also highly expressed in undifferentiated spermatogonia stem cells and their somatic support cells in testis [32] (our unpublished data). Loss of PTBP1 in spermatogonia cells disrupts spermatogenesis and induces apoptosis of these cells [32]. Loss of PTBP1 in Sertoli cells impairs the supporting role of Sertoli cells, leading to failure of spermatogenesis (our unpublished data). Thus, it is tempting to speculate that, in tissues that undergo constant regeneration, PTBP1-mediated post transcriptional regulation is essential for maintaining stem cells and progenitor cells multipotent and viable.

Results from our global transcriptome analysis show that, in addition to *Phlda3* and *Ptbp2*, PTBP1 controls alternative splicing of many other genes in the crypt cells. Loss of PTBP1 causes both an increase and a decrease in exon inclusion in the crypt cells. The increased exon inclusion events, however, are significantly more than the decreased events, suggesting that PTBP1 functions primarily as a repressor of alternative spliced exons in the crypt cells. This is consistent with previous finding that PTBP1 either represses or promotes exon inclusion of alternative exons depending on its binding sites in pre-mRNA [25]. Using exon ontology, we further show that PTBP1-regulated alternative exons are important for protein structure, localization, modification, binding with other proteins, and catalytic activity. Intriguingly, we found these exons are significantly enriched for regions encoding phosphorylation sites, suggesting that PTBP1 regulates the functions of its target genes through the control of their protein phosphorylation. Through gene ontology, we show that PTBP1-mediated splicing control is important for biological processes that modify gene function at the mRNA and protein levels, such as the splicing control, phosphorylation, methylation, and protein binding. While our findings support that upregulation of *Phlda3* causes crypt cell death in PTBP1-deficient mice, we cannot exclude the possibility that the aberrant splicing of other genes induced by PTBP1 deficiency contribute to crypt cell death as well.

Our studies using the constitutive *Villin* promoter linked-*Cre* mouse model and the tamoxifen-inducible *Cre* mouse model for PTBP1 deletion indicate that PTBP1 plays age-related roles in IECs [34]. In the neonatal stage, we show that PTBP1 deletion mediated by the constitutive *Villin* promoter linked-*Cre* recombinase does not result in epithelial regeneration defects in the small intestine [34]. Instead, the knockout mice develop intestinal inflammation in the colon shortly after birth, which is accompanied by early onset of colitis and colorectal cancer [34]. Mechanistically, we demonstrate that epithelial PTBP1 downregulates the Toll-like receptor signaling activity to suppress the intestinal immunity in the neonates, which is important for generating a permissible environment for gut microbiota formation [34]. The discrepancy in PTBP1 functions in neonatal stage and adulthood are likely due to the difference in epithelial regeneration mechanisms between the two stages. Unlike the adult small intestine wherein Paneth cells support ISC survival and proliferation, the neonatal intestine is immature and Paneth cells do not appear in the neonatal intestine until the second week after birth [45, 46]. As such, a Paneth cell-independent mechanism accounts for the ISC survival and regeneration in the early neonatal stage [47]. In addition, the neonatal stage is a critical time window for microbial colonization which requires transient intestinal immune suppression [48]. As such, the main physiological role of PTBP1 in the neonatal intestine is presumably to mediate immune suppression instead of regulating ISC survival and proliferation.

## Materials and methods

### Animals

#### Generation of *Ptbp1*^flox/flox^; *Villin-cre*^ERT2/+^ mice

The floxed *Ptbp1* mouse allele *Ptbp1*^flox/flox^ (hereafter *Ptbp1*^f/f^) was generated as previously reported [34]. The floxed *Ptbp1* mice were crossed with the *Villin-creERT2* mice [49] to generate the *Ptbp1*^flox/flox^; *Villin-Cre*^ERT2/+^ mice (hereafter *Ptbp1*^f/f^; *Vil-cre*^ER+/−^). Primers used for genotyping the *Ptbp1* floxed allele were forward: 5’–CCCATAACTGTCCATAGACC-3’, and reverse: 5’-TGTTGGTAATGCCAGCACAG-3’. Primers used for genotyping the *Villin-creERT2* in these mice were: Forward: 5’-ATGTCCAATTTACTGACCGTACACC-3’, and Reverse: 5’-CGCCTGAAGATATAGAAGATAATCG-3’.

All mice used in this report are from the cross of the *Ptbp1*^f/f^; *Vil-cre*^ER+/−^ mice with the *Ptbp1*^f/f^ mice. The tamoxifen administrated *Ptbp1*^f/f^; *Vil-cre*^ER+/−^ mice are referred to as the knockout mice, and their littermates *Ptbp1*^f/f^ mice administrated with the same dose of tamoxifen are referred to as the control mice. Tamoxifen was administrated by single daily injections for two consecutive days at the dose of 100 mg/kg of body weight.

#### Generation of *Ptbp1*^flox/flox^;*Vil-cre*^ERT2+/−;^*Lgr5-Egfp-IRES-creERT2*

*Ptbp1*^f/f^;*Vil-cre*^ER+/−^ mice were crossed with *Lgr5-Egfp-IRES-creERT2* mice (the Jackson Laboratory) to generate *Ptbp1*^flox/flox^;*Vil-cre*^ERT2+/−^; *Lgr5-Egfp-IRES-creERT2* (hereafter *Ptbp1*^f/f^;*Vil-cre*^ER+/−^;*Lgr5*). Primer used for genotyping the *Villin-creERT2* in these mice are forward: 5’-GTGTGGGACAGAGAACAAACCG-3’, and reverse: 5’-TGCGAACCTCATCACTCGTTGC-3’. *Lgr5-Egfp-IRES-creERT2* was detected as described by the Jackson Laboratory. *Ptbp1*^flox/flox^;*Lgr5-Egfp-IRES-creERT2* (hereafter *Ptbp1*^f/f^;*Lgr5*) mice were used as the control mice.

#### Weight loss measurement and death record

Mouse body weight was measured before tamoxifen injection and every 24 hours after tamoxifen injection for 5 consecutive days. Percent weight loss was calculated by subtracting the weight measured at 24, 48, 72 hours, etc., post tamoxifen injection from the weight measured at 0-hour post tamoxifen injection and dividing by 0-hour weight. Mice were monitored every day after tamoxifen injection for at least two weeks.

#### Histology and Immunostaining

Intestines were isolated, fixed in 4% paraformaldehyde at 4□ overnight, paraffin-embedded, and sectioned according to the standard protocols. Intestine sections (5 μm) were processed for hematoxylin and eosin staining or for immunostaining. Immunohistochemistry was performed using the R.T.U. vectastain kit (Vector Laboratories) with DAB substrate and sections were counterstained lightly with hematoxylin afterwards. Double immunofluorescence staining with two antibodies produced in rabbits was done by following a protocol described at the Jackson Immune Research Laboratory website (https://www.jacksonimmuno.com). Briefly, after the first secondary antibody incubation, rabbit serum was applied to saturate open binding sites on the first secondary antibody. This was followed by applying excessive unconjugated fab goat anti-rabbit IgG (H+L) fragments on the sections to cover the rabbit IgG to prevent the binding of the second secondary antibody to the first primary antibody.

Secondary antibodies used are goat anti-rabbit AlexaFluor 488 or 594, goat anti-mouse AlexaFluor 594, and donkey anti-goat AlexaFluor 488 (all from Invitrogen). Sections were counterstained with 4’,6-diamidino-2-phenylindole (DAPI). Primary antibodies used are rabbit anti-active CASPASE 3 (Cell Signaling, 9661), mouse anti-KI67 (BD Pharmingen, 550609), rabbit anti-OLFM4 (Cell Signaling, 39141), goat anti-PTBP1 (Santa Cruz sc-16547), or rabbit anti-PTBP1 (gift from Dr. Douglas Black), goat anti-GFP (Rockland, 600-101-215), rabbit anti-P53 (Leica Biosystems, NCL-L-p53-CM5p), and rabbit anti-PHLDA3 (LifeSpan BioSciences, LS-C499531-100).

Images were taken from a Leica compound microscope with a digital camera or a Nikon A1R confocal microscope and processed using Adobe Photoshop.

#### Intestinal organoid culture and whole mount immunostaining

Small intestinal crypts were isolated from *Ptbp1*^f/f^; *Vil-cre*^ER+/−^ and *Ptbp1*^f/f^ mice as described [50] and cultured in IntestiCult Organoid Growth Medium (Mouse) (StemCell Technology, 06005). 4-hydroxytamoxifen was added into the culture medium at a final concentration of 200 nM to delete *Ptbp1*. After 24 hours incubation, 4-hydroxytamoxifen was removed by replacing with fresh medium. Culture medium was changed every 48 hours.

For whole mount immunostaining, organoids were fixed in 4% paraformaldehyde at 4□ overnight, washed with phosphate buffered saline (PBS) and permeabilized with 0.5% Triton x-100 in PBS. Organoids were then treated with 100 mM glycine in PBS to block free aldehyde groups. After being washed with PBS, organoids were treated with PBS containing 5% serum, 0.1% BSA, 0.2% Triton x-100, and 0.05% Tween for 90 minutes at room temperature to block nonspecific antigen binding and incubated with the primary antibody overnight at 4□. After washed with PBS, organoids were incubated with the secondary antibody in the dark at room temperature for 2 hours and counterstained with DAPI. Images were taken from a Nikon A1R confocal microscope and processed using Adobe Photoshop.

#### Quantitative Real-Time PCR

Small intestinal crypt cells were isolated as described [50] at 20, 24, 36 and 48 hours post tamoxifen induction. Briefly, small intestines were harvested and flushed with ice-cold PBS. The intestines were cut into ~5 cm fragments and open longitudinally and washed with ice-cold PBS. After villi were gently scraped off, fragments were incubated in 2 mM EDTA/PBS with gentle shaking at 4°C for 10 min and pipetted up and down 10 times with a transfer pipette. After the tissues were settled down, the supernatant was removed. Intestine fragments were then incubated in 2 mM EDTA/PBS with gentle shaking at 4°C for additional 20 minutes. After the supernatant was removed, fragments were washed with cold PBS and vortexed for 30 seconds with 3 seconds pulses. The fragments were allowed to settle, and the supernatant was collected and set aside on ice. This process was repeated twice. The collected supernatant fractions were filtered through 70 μm pore cell strainers (BD Biosciences, Bedford, MA). The filtrate was centrifuged at 200 *g* for 5 min, and the pellet was used for RNA extraction or western blots. RNAs were extracted using TRIzol reagent according to standard protocols. Real-time PCR reactions were performed blindly in triplicate or duplicate using SYBR green master mix. PCR primers used are listed in supplemental table 1.

#### Western Blots

Isolated small intestinal crypt cells were homogenized in lysis buffer as described [34]. Protein lysates were cleared by spinning the samples twice at 4°C. Subsequently, samples were separated on SDS-PAGE and analyzed by western blotting as described [34]. Primary antibodies used are mouse anti-PTBP1 (Life Technologies, 324800), mouse anti-PTBP2 antibody (Santa Cruz, sc-376316), rabbit anti-Pan AKT (Cell Signaling, 4685), rabbit anti-Phospho-AKT (Ser473) (Cell Signaling, 4060), rabbit anti-GAPDH (Santa Cruz Biotechnology, SC-32233), and rabbit anti-GRP (Abcam, ab32618). Membranes were incubated with HRP-linked secondary antibodies and developed using ECL prime (G&E Healthcare Life Sciences).

#### RNA-seq analysis

Crypt cells from small intestines were isolated at 20 hours post tamoxifen administration from 3 knockout mice and 3 littermate control mice. RNAs were extracted using PureLink RNA Mini Kit (Ambion, Cat. 12183025). RNA-seq libraries were constructed and sequenced at the UIUC Biotechnology Center High-Throughput Sequencing Core. Sequencing was done by using 150 base pairs paired-end reads. Each library generated over 1.7 million reads. Raw reads were subjected to read length and quality filtering using Trimmomatic V0.38 [PMID: 24695404] and aligned to the mouse genome (mm10) using STAR (version 2.6.1d) [PMID: 23104886]. Cufflinks package [PMID: 22383036] was used to assess differential gene expression events, among which significant events were identified using a stringent cutoff criteria (FDR(q-value) < 0.05, log2(fold change) > 1). rMATS v4.0.2(turbo) [PMID: 25480548] was used to study differential splicing, and events with FDR < 0.1, junction read counts ≥ 10, PSI ≥ 10% were deemed to be significant.

Exon ontology analysis was performed on the set of alternatively spliced cassette exons identified using rMATS. Mouse(mm10) annotations were converted to human (hg19) annotations using UCSC liftover with a minimum base remap ratio set to 0.8. Exon ontology pipeline [PMID: 28420690] was then used on the lifted exons to perform ontology analysis.

#### Statistics

Differences between the knockout mice and the control groups were assessed for significance using a two-tailed unpaired Student t-test. Data involving two or more variables were analyzed by two-way ANOVA using GraphPad Prism.

## Data availability

All raw RNA-seq data files are available for download from the Gene Expression Omnibus (accession number GSE185499).

## Ethics statement of animal use

All procedures involving mouse care, euthanasia, and tissue collection have been approved by the University of Illinois at Urbana-Champaign Animal Care and Use Committee (IACUC approved protocol #20177 and 20211). Mice were used and cared for according to the institutional “Guide for the Care and Use of Laboratory Animals” and in accordance with all University of Illinois at Urbana-Champaign policies and guidelines outlining the care and use of animals in research.

## Acknowledgments

We thank Dr. Adam Snook and Dr. Sylvie Robine for providing the villin-creERT2 mouse line, Dr. Douglas Black for providing anti-PTBP1 antibody, Dr. Waqar Arif for sharing computational tools, and Dr. Bo Wang for the technical support on organoid culture.

## Author contributions

W.S.T designed the project, performed the experiments, interpreted the data. V.C.U. performed computational analysis and the experiments, interpreted the data, helped writing the manuscript. K.L.N, A.L., D.Y., K.C., and J.Y. performed experiments. S.B. assisted in performing computational analysis. W.M. designed the project, performed the experiments, interpreted the data, supervised the study, and wrote the manuscript. A.K designed experiments, interpreted the data, supervised the study, and helped writing the manuscript.

