## Supplementary figures and images for "PTBP1/HNRNP I controls intestinal epithelial cell regeneration by maintaining stem cell survival and stemness"

### Supplemental data

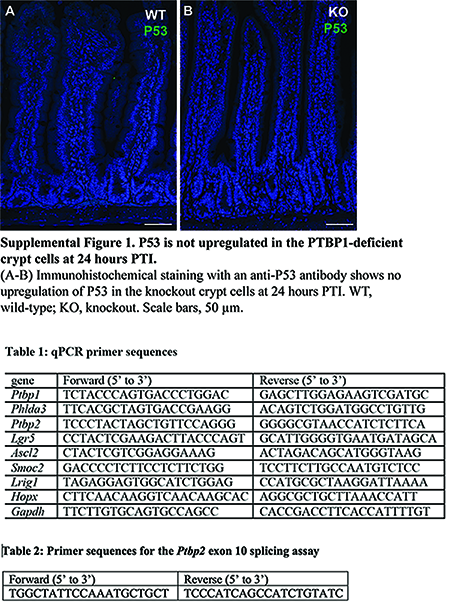
